# CellPheno: A High-throughput Computational Platform for Quantifying Cellular Resolution Whole Brain Microscopy Images

**DOI:** 10.64898/2026.03.17.712391

**Authors:** Ziquan Wei, Ian Curtin, Felix A Kyere, David Borland, Hong Yi, Minjeong Kim, Mustafa Dere, Carolyn M McCormick, Oleh Krupa, Yen-Yu Ian Shih, Mark J Zylka, Jason L Stein, Guorong Wu

## Abstract

Advances in tissue clearing and light-sheet microscopy enable cellular resolution whole-brain 3D imaging. However, whole-brain quantification tools do not yet meet demands for efficiency or assess morphometry. Here we present *CellPheno*, a 3D nuclei instance segmentation framework for high-throughput cellular phenotyping. *CellPheno* quantifies an entire P4 mouse brain within 15 hours. We showcase whole-brain morphometry, enhanced stitching, and co-localization across multiple cell types in 53 brains.

## Main text

Our understanding of nervous system function is critically dependent on visualizing and quantifying the three-dimensional (3D) structure of the brain. Recent innovations in tissue clearing techniques and light-sheet fluorescence microscopy (LSFM) enable us to image brain samples with 3D micron resolution to quantify cellular phenotypic differences in brain tissue^1,2^. Although the counting and localization of whole brain cells in the real world has been conducted on mouse^3^ and fish^4^ models, there is currently no cellular phenotyping platform to investigate morphological characteristics through 3D nuclei instance segmentation (NIS). Morphometric features derived from NIS, however, have been shown to be more informative than cell localization alone for understanding brain aging and disease, such as nuclear circularity in amyloid plaque from mouse brains with Alzheimer’s disease (AD)^5^, dysmorphic nuclei from rat brains with autism spectrum disorder^6^, human stem cells with frontotemporal dementia^7^, and Drosophila brains expressing human tau^8^. Several nuclei segmentation methods have been developed^9–12^, however current approaches lack scalability for whole-brain instance segmentation involving ∼100 million cells. Specifically, existing methods are designed for relatively small image patches, hindering their scalability for whole-brain analysis.

As microscope and image stitching technologies advance, it is likely that in the near future cellular resolution maps of the entire adult human brain containing 170 billion cells (86 billion neurons) can be produced, significantly increasing the demand for quantitative platforms to analyze whole brain nuclei at cellular resolution^9,13^. In parallel, quantitative analysis of brain structures from LSFM enables us to measure and compare regional variations such as nuclei shapes and volumes, with high precision. Nonetheless, compared to the numerous analytics pipelines for human brain neuroimaging^14,15^, the cellular computational platform for large-scale LSFM of mouse brains is still limited to cell counting and localization only^16–19^. To meet these emerging needs, there is a strong demand for a computationally efficient and scalable NIS pipeline that produces consistent readouts of nuclei morphometry across neuroscience studies to benefit downstream brain-wide analysis.

In this work we present a high-throughput computational platform of cellular phenotyping (*CellPheno*) for analyzing real-world whole-brain microscopy images with multiple labels. In contrast to the progress made in 2D whole slide image NIS^20–22^, systematic and resolution-independent NIS pipelines in 3D remain scarce. Although the emerging associations^5–7^ between 2D nuclear morphology and brain pathology make full 3D volumetric measurement indispensable, directly extending 2D approaches to 3D lose accuracy due to the anisotropic resolution in LSFM^23^. Furthermore, acquiring and processing these massive 3D brain volumes with limited CPU memory necessitates imaging and computing in smaller, overlapping tiles (**Fig. 1 a**), resulting in the major challenge of stitching tiles and reconnecting fragmented cells located at tile boundaries. *CellPheno* is designed as a 2D-to-3D interactive framework that features accuracy, efficiency, anisotropic resolution handling, and accessibility of local and global quality control (**Fig. 1 b**). We first utilize Cellpose to predict accurate 2D probability and spatial gradient maps (**Fig. 1 c**). *CellPheno* then efficiently rebuilds only the spatial gradient along the third dimension from 2D using a median filter pyramid (**Fig. 1 d**), where the 3D boundary can be extracted (**Fig. 1 e**). Afterward, stitching in the cropped (**Fig. 1 f**) and overlap area (**Fig. 1 g**) is achieved by a graph neural network (GNN) and the point registration between NIS, respectively, omitting convolutional kernels that require isotropic resolution. Lastly, the extraction of a brain representation map of various morphological features (**Fig. 1 ij**) can be easily accessed by averaging nuclei instances within every cubic area (**Fig. 1 k**) for downstream neuroscience analysis (**Fig. 1 l**) and applications (**Fig. 1 m**).

**Figure 1:**
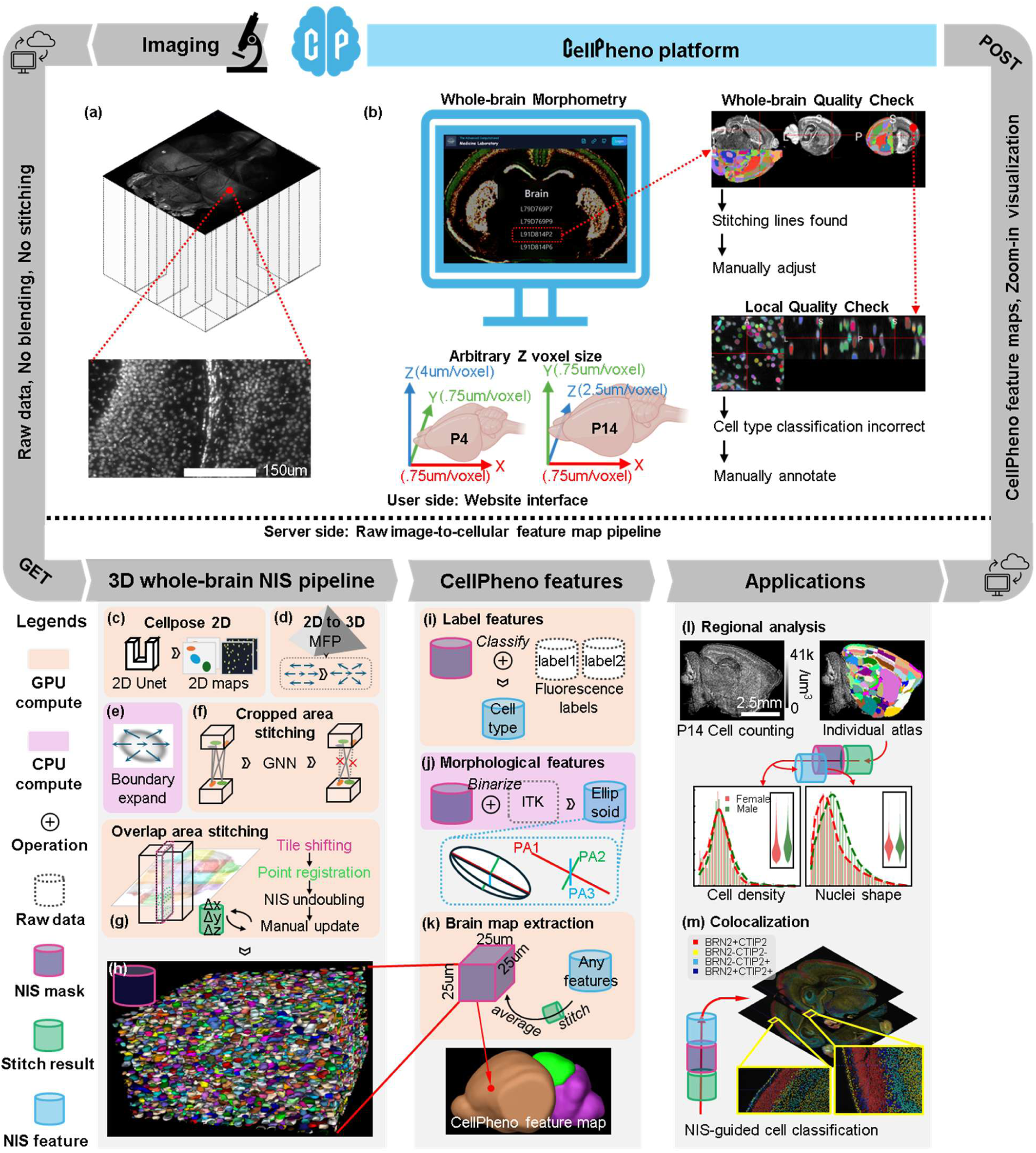
A high-throughput, quality-control enabled computational platform for cellular phenotyping to extract morphological features from whole-brain nuclear resolution images. a: Teravoxel whole brain microscopy images are imaged by overlapping tiles. *CellPheno* processes the tiles instead of stitched images to eliminate the accumulation of errors during preprocessing. b: *CellPheno* can handle arbitrary Z resolutions and enables users to control the quality of whole-brain stitching and local classification. c: *CellPheno* segments cell instances based on 2D predictions from Cellpose^10^. d: 3D spatial gradient maps are rebuilt by a 2D-to-3D approach: median-filter pyramid (MFP). This 2D-to-3D approach makes *CellPheno* high-throughput. e: 3D instance masks are constructed in the same manner as in Cellpose^10^ by using 3D probability maps stacked from 2D maps and the 3D gradient maps from MFP. f: Cropped area stitching is achieved using a Graph Neural Network (GNN). g: Overlapping area stitching is achieved by the coarse-to-refine workflow. h: Examples of NIS masks within a local 25μm^3^ cube. i: The multi-channel fluorescence label features are used to determine the cell type. j: The binarized NIS mask is processed by ITK^24^ to obtain the principal axes (PAs). k: *CellPheno* whole-brain feature map extraction. l: The individual registration from atlas space to *CellPheno* brain map leads to regional analysis. m: Colocalization is guided by NIS results to focus only on the nuclei area.

### Computational time

On a local rack server equipped with 128 CPU cores and a 48GB GPU, *CellPheno* processed each mouse brain in 15 hours, operating in less than half the time of three existing whole-brain cellular NIS approaches^10,16,18^ while maintaining performance above 0.9 in both recall and precision. The NIS accuracy from *CellPheno* generally outperforms Cellpose 3D 1.0 ^10^ (**Fig. 2 a**) (93.0 to 89.8 in F1). The running time distributes linearly from 13 to 18 hours according to tissue volume from 50 to 75 mm^3^ (**Fig. 2 b**), whereas other pipelines representing cells as points cannot process a single brain brain in one day to meet the LSFM imaging speed, i.e., 20∼30 hours per brain ^16^.

**Figure 2:**
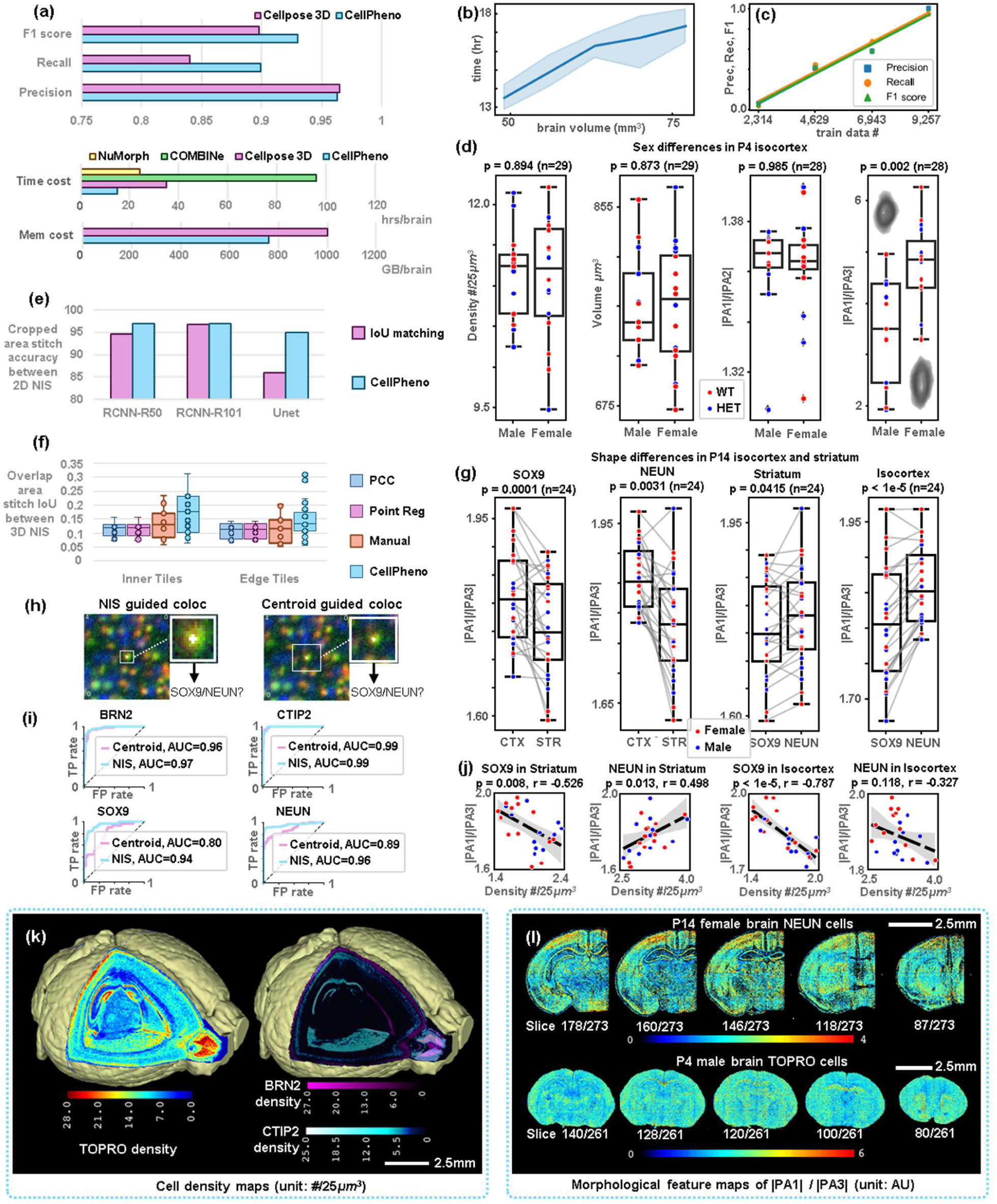
Whole-brain NIS and co-localization performance, statistics, and readout. a: Segmentation accuracy and computational costs of *CellPheno*. b: Speed of *CellPheno* w.r.t. microscopy image size, tested on thirty P4 mouse brains. c: Sample size of manually annotated 3D nuclear segmentations (training data) versus segmentation precision, recall, and F1 scores using 0.3 IoU (Intersection over Union) threshold. d: Independent *t*-tests between male and female mice for different morphological features derived from *CellPheno*, where two gray models visualize the average NIS shapes of male and female. e: NIS stitching accuracy in the cropped area between the conventional IoU matching and *CellPheno*. f: IoU between NIS in the overlapping area after stitching indicates the image stitching accuracy. Phase correlation correction (PCC) is coarse stitching; point registration (reg) is refined stitching; and manual is performed by IC and FAK using the Imaris Stitcher. g: Paired *t*-tests show spatial and cell type differences in Isocortex (CTX) and Striatum (STR) regions. h: NIS-guided co-localization uses the precise nuclei area as input. In contrast,centroid-guided uses a fixed window. i: Area under curve (AUC) of Receiver Operating Characteristic (ROC) by NIS or Centroid guided co-localization. j: The correlation between cell density and nuclei shape in P14 mice (n=24), where each dot is a mouse. k: *CellPheno* whole-brain density maps of TOPRO, BRN2, and CTIP2 cells in P4 brain. l: Multiple coronal slices of whole-brain morphological feature maps from P4 and P14 brains.

### Accuracy of nuclei instance segmentation

We have evaluated the NIS accuracy with respect to the size of the labeled training dataset and found that the annotated training data used in *CellPheno* is sufficient to achieve high precision, recall, and F1 scores (**Fig. 2 c**).

### Applications

By capitalizing on the computational power of identifying each nucleus in the entire brain image, *CellPheno* enables a range of novel applications in neuroscience research.

● *Shape analysis.* Morphological analysis of nuclei shapes is possible after identifying each nucleus instance by *CellPheno*, where significant shape differences in terms of the length ratio between principal axes |PA1|/|PA3| were detected between female and male mice, at the significance level p value=0.002, n=28 (**Fig. 2 d**). In contrast, neither cell counting nor volume yielded a significant sex difference.
● *Cell co-localization* based on multi-channel LSFM, i.e., *BRN2*/*CTIP2* and *SOX9/NEUN*, in addition to *TOPRO*, which *CellPheno* segments on, is achieved using NIS-guided CNN (**Extended Fig. 7**). Due to precise patch cropping guided by the NIS against the centroid (**Fig. 2 h**), NIS enabled superior classification performance (**Fig. 2 i**).
● *Brain mapping of nuclei development.* Given the NIS results, we summarize nuclei density in every 25×25×25μm^3^ cube in the brain and yield a brain mapping of nuclei development

(**Fig. 2 kl** and **Extended Fig. 9**), which can be treated as a ready-to-use representative mapping of brain structure.

*CellPheno* exhibits high efficiency for whole-brain 3D nuclei instance segmentation and scalability for emerging applications in spatial cell phenotyping. Together with the web-based computational platform, *CellPheno* enables efficient processing, visualization, and quantitative analysis of whole-brain cellular data, which opens new opportunities for region-specific characterization of brain structure and facilitates research in neurodevelopment and neurodegeneration.

## Methods

### Overall pipeline

*CellPheno* is a computationally efficient, robust, and scalable whole-brain cellular phenotyping framework that incorporates advancements in cellular resolution and standardization enabled by LSFM imaging and 3D NIS. To achieve optimal acceleration, it is recommended to deploy CellPheno in a computing environment equipped with 2×48GB GPU memory and 1TB of CPU memory. As shown in **Fig. 1 c-k**, CellPheno performs tensor calculations on the GPU and iterative calculations on the CPU, with GPU and CPU computations executed in parallel for efficiency. The project’s code and documentation are version-controlled and available on <u>GitHub</u>.

*CellPheno* extends the idea of Cellpose to 3D NIS. It first predicts the probability of nuclei and their spatial gradients on 2D planes with high isotropic resolution, i.e., 0.75 μm per voxel. A median filter pyramid (MFP) is then used to reconstruct the spatial gradient along the third dimension (typically 2.5 μm or 4 μm per voxel), achieving a balance between efficiency and accuracy in 3D NIS. All components of CellPheno are quality-control enabled and parallelizable.

### Tissue Clearing and Immunolabeling

We have applied *CellPheno* to an imaging-genetics study^25^ involving 29 P4 mice (M/F: 17/12), which are stained with three dyes: all nuclei by TO-PRO-3 (TOPRO), upper-layer neurons (Layers II/III) by the BRN2 marker, and deeper-layer neurons (Layers Vb/VI) by the CTIP2 marker. 24 P14 (M/F: 13/11) mice with glial and neuronal cells were also stained by SOX9 and NEUN, respectively. All 53 samples were batch processed in littermate pairs. All samples were imaged intact on a light-sheet microscope within 1 week of completing tissue clearing and immunolabeling. For further details refer to the study^25^.

### Manual Annotations

We used a stand-alone application, Segmentor^26^, and its web-based extension enabling crowdsourced nuclei annotation, ninjato (https://github.com/renci/ninjato), to annotate the full 3D shape of 10,827 nuclei in lightsheet images of the mouse brain at both P4 and P14. In total, 24,463 2D patch images were annotated with cell type labels. The number of samples per class is as follows: 9805 Brn2-Ctip2-, 4400 Brn2+Ctip2-, 3000 Brn2-Ctip2+, 1030 Brn2+Ctip2+, 1960 NEUN-SOX9-, 868 NEUN+SOX9-, 3251 NEUN-SOX9+, and 149 NEUN+SOX9+.

### Standard input

The raw volumetric image from the LSFM scanner is commonly stored as multiple 2D slice images. CellPheno directly takes 2D slices as input. Intensity of each is first normalized to the range from 0 to 1 after dropping 1% lowest and highest intensities. Then, the 2D slices are resampled to the same resolution as the training using bilinear interpolation. Last, the 2D slices are cropped to 224×224 patches with 10% overlap. Each of the four hundred patches are batched as one input of the following U-shaped convolutional neural networks (Unet) (**Extended Fig. 1**).

### Training

We train Unet using all 2D slices extracted from the 9,257 3D nuclei annotations. We have evaluated NIS accuracy on 1,570 manually annotated 3D nuclear nuclei. The output of Unet after patching is a 3-channel 2D map: voxel probability as a nucleus and two spatial gradients in the horizontal and vertical directions. Training is performed by supervised learning between the output and the annotation generated from the 3D segmentation ground truth. First, for the probability map, we convert each 3D segmentation mask into a 2D binarized map as the foreground annotation, supervising the probability map via cross-entropy. Then, for the spatial gradients, we compute the shortest distance from the voxel to the instance segmentation boundary, so that the gradient points toward the pseudo-center of the nucleus. Unet was trained for 100 epochs with stochastic gradient descent (SGD) with a learning rate of 0.2, a momentum of 0.9, a batch size of eight images, and a weight decay of 0.00001. The learning rate was initialized at 0 and linearly annealed to 0.2 over the first 10 epochs to prevent initial instabilities. For all analyses in the paper, we used a base of 32 feature maps in the first layer, increasing by a factor of 2 at each downsampling step, reaching 256 feature maps in the layer with the lowest resolution.

### MFP: 2D-to-3D approach

First, the prediction for the third dimension is rebuilt using a median filter pyramid (MFP, Extended Fig. 2), a dynamic estimator that computes the L2 norm of the 2D gradient between the previous and next slices. We initialize the third gradient as the next-slice gradient norm minus the previous-slice norm. For voxels with similar norms within the nucleus, we iteratively update the norm map using a 3×3 median kernel, and average the differences between the filtered norms from each iteration as the third gradient. This enables nuclei of varying sizes to be gradually covered by repeating the iteration. We then threshold the probability map and stack the 2D maps along the third dimension to obtain 3D maps, retaining only probabilities greater than zero. A dynamic system starts at every foreground voxel and follows the spatial derivatives specified by the 3D gradient. At each iteration, a 2-voxel-long step was taken towards the direction of the gradient at the nearest foreground grid location. Following convergence, voxels are clustered by their resulting locations. For robustness, the clusters are extended by 3 voxels along regions of high-density foreground convergence to prevent the exact center from being missed by Unet in certain cases.

### Model readout

The standard output is a collection of 3D coordinates, denoted as {**P** ∈ ℝ^*V*×3^}, of all voxels in each cluster, where N is the number of instances, and V refers to the volume size of a nucleus instance. This standardization benefits downstream applications and morphometric analysis, which are applied independently for each instance. Meanwhile, the center of each instance is stored before downstream analysis as the world coordinate for *CellPheno* feature generation, as shown in **Fig. 1 k.** The whole-brain density map therefore provides a whole-brain visualization rather than requiring stitching and blending of LSFM images.

### Stitch cropped NIS masks

For the output coordinates located at the cropping boundary of the 3D input, due to finite CPU memory, we construct a graph from the cropped NIS masks and train graph neural networks to stitch them, as shown in **Fig. 1 b**. The coordinates at the cropping boundary of each cluster are first collected as nodes in the graph, and each node is assigned a 227-dimensional vector as its feature. The first 200 are the resampled image intensities within the NIS mask, and the last 27 are the counts of 27 possible directions of the 3D spatial gradient (**Extended Fig. 3 c**). Edges of the graph are connected to the top 2 closest nodes and are assigned with the spatial distance between centers, height and width ratios, and the average spatial gradient (**Extended Fig. 3 b**). Graph convolution networks (GCN, **Extended Fig. 3 e**) take this graph as input and classify if the edges connect the same nuclei as the output. GCN training is supervised using a cross-entropy loss between the output and the annotation generated from the 3D segmentation mask. Every two neighboring slices of the 3D mask were used to generate the annotation. GCN was trained for 300 epochs with SGD with a learning rate of 0.001, a momentum of 0.9, a batch size of 64 pairs of neighboring mask slices, and a weight decay of 0.00001. The learning rate started from 0.001 and was linearly annealed with a weight of 0.5 every 30 epochs.

### Stitch the overlapping area of tiles

For the output coordinates located at the overlapping area of two neighboring tiles, there are two stages to prevent double segmenting: (1) a 3D Inter-over-Union ratio (IoU) guided wobbly stitching^27^ by non-rigid registration and (2) undoubling the segmentation by IoU matching. The wobbly stitching starts from a phase correlation correction (PCC) between every pair of 3D tiles to estimate a coarse stitching transformation for the entire 3D tile, and then applies the average transformation from all neighbors to each 3D tile. The coarse stitching has to load whole-brain LSFM images, resulting in 2 hours of computational time. Next, the point registration is applied as refined stitching to every 25 slices between a coarsely stitched tile and all neighboring refined tiles (**Extended Fig. 4**), where points are represented by the center of NIS.

### Co-localization

The co-localization on multiple channels, e.g., BRN2 and CTIP2, in addition to TOPRO3, which CellPheno segments on, is achieved using NIS-guided ResNet50. Considering the unbalanced SNRs between different channels, the global location of a cell is an important prior knowledge for the manual annotation. We implement a Matlab annotation tool to display 1,000 local patches of NIS results alongside a whole-brain location indicator (**Extended Fig. 5 ab**). Local patches can be randomly picked or explicitly selected from interesting regions with a brain mask. Neuroscience experts determined the patch label based on the information of multichannel intensity and whole-brain location. For either training or inference, the patch image of each channel is first normalized to [0,1], and then stacked as a multichannel LSFM image, cropped by the NIS bounding box, and resized to 30×30. During the training (**Extended Fig. 5 c**), there are three image augmentations randomly (50%) applied to the image for training data variability. Additionally, (i) the spatial location (x,y,z) is embedded and concatenated to the latent feature from the patch, and (ii) the imbalanced sampling applied according to the cell count per channel is uneven across the whole brain. The training of ResNet50 ^28^ has 500 epochs and a 1e-4 learning rate using the Adam optimizer. For inference (**Extended Fig. 5 d**), the image augmentations were skipped. For P4 BRN2/CTIP2 multichannel LSFM images, our annotations were selected in subregions of the isocortex where upper and lower layer neurons mainly existed (**Extended Fig. 5 d**). Note that the slice index is along the dorsal-ventral direction.

### Morphometry analysis

Morphometry of the NIS mask measures the ellipsoid of the nuclei (**Extended Fig. 6**). First, the 3D boundary of each NIS mask is extracted using the Open Computer Vision (OpenCV) library. Then, the endpoints of three Feret diameters are determined among the boundary points using the Simple Insight Toolkit (SimpleITK^24^). Finally, the nucleus ellipsoid is indicated by |PA1| / |PA2| and |PA1| / |PA3|, where PA1, PA2, and PA3 are the Feret diameters along the three principal axes.

### Statistical analysis

The Allen P4 mouse brain atlas is first registered to each individual using Advanced Normalization Tools (ANTs) with the LSFM image downsampled to 25μm/voxel (**Extended Fig. 7**). An independent t-test is then computed between the isocortex region of male and female mice, such that the most interesting regions are statistically analyzed. For every CellPheno measurement, samples with a z score greater than 3 are treated as outliers and removed during the statistical analysis.

### Website interface

We developed a web interface for CellPheno to analyze nuclei from LSFM images. The computation of NIS runs on the server side, followed by an email notification to users when NIS has been completed and whole-brain results have been successfully uploaded back to the Amazon cloud. The whole-brain result includes all available feature maps (25μm/voxel) along with instance segmentation masks at the raw resolution saved in small chunks (125×125×75μm^3^). They are accessible in the whole-brain view and the zoom-in view, respectively (**Extended Fig. 8**), where users can browse and perform quality control. All volumetric data in the browser page can be downloaded for downstream analysis.

## Data Availability

The raw multi-channel light sheet images supporting the results of this study have been deposited in the Brain Observatory Storage Service and Database (BossDB) and are publicly available at https://bossdb.org/project/curtin2026.

## Code Availability

*Cellpheno* code isavailable at https://github.com/Chrisa142857/Lightsheet_microscopy_image_3D_nuclei_instance_segmentation.

---

**Extended Fig. 1:**
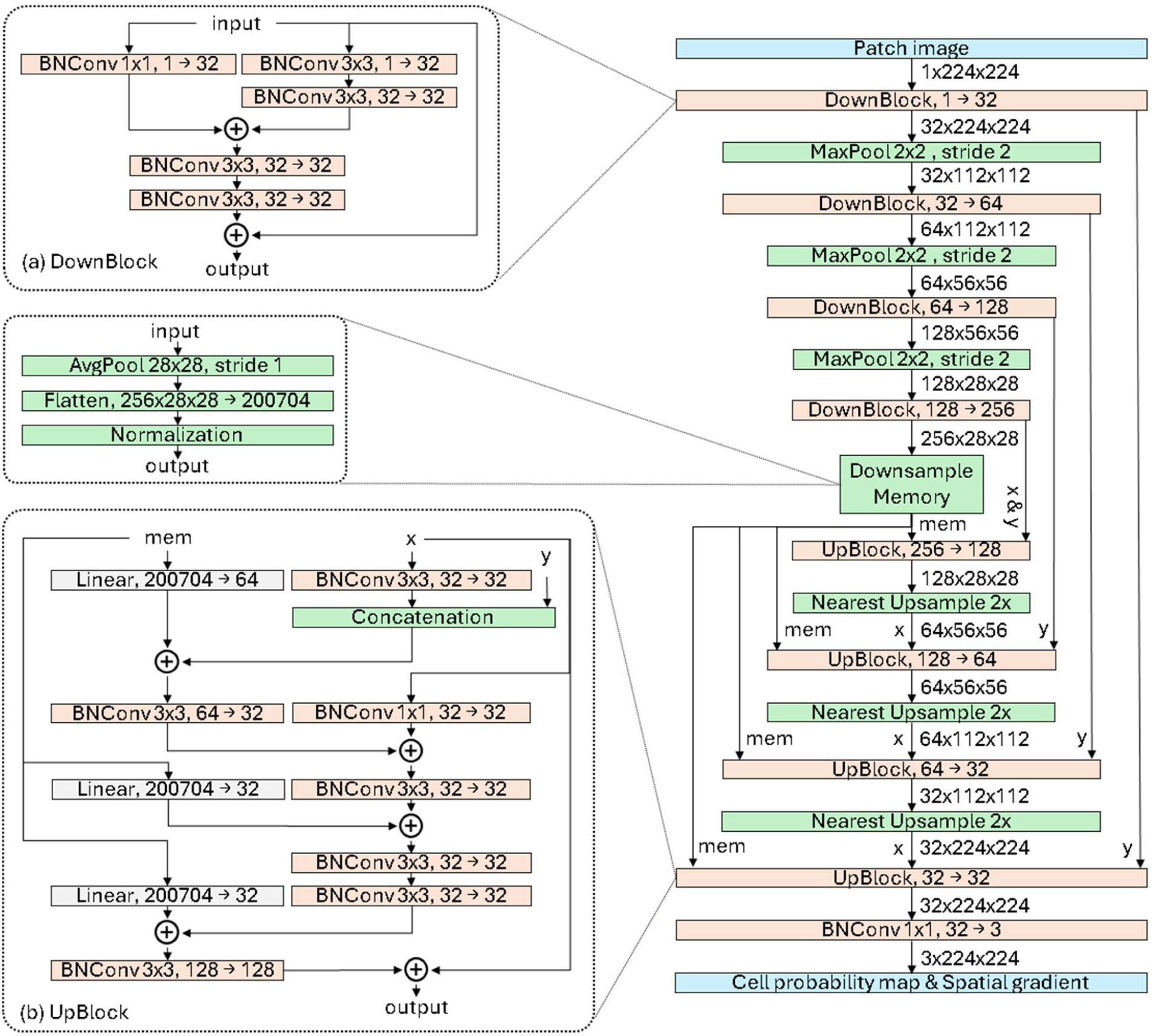
Unet architecture. a: Downsampling block, where ‘BNConv’ denotes one convolution layer following the batch normalization and the ReLU activation. b: Upsampling block, where ‘Linear’ denotes one layer of linear transformation using <u>torch.nn.Linear</u>.

**Extended Fig. 2:**
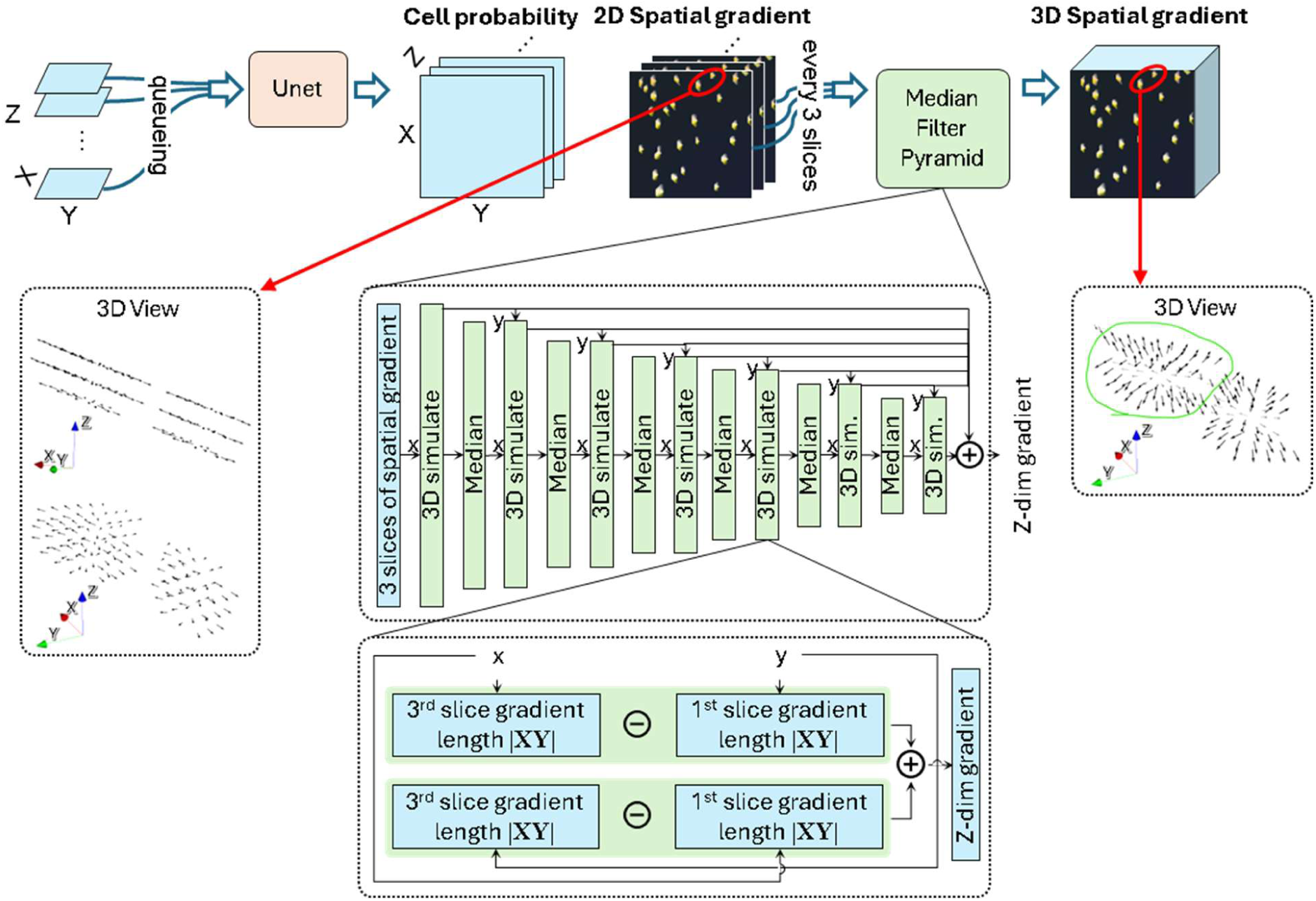
MFP architecture. Flowchart of predicting the third dimension (Z-dim) spatial gradient, where ‘Median’ denotes one median filter pooling layer with a 3×3 kernel size, which is implemented by kornia.filters.median_blur, and |**XY**| denotes the length of the gradient vector in the xy plane.

**Extended Fig. 3:**
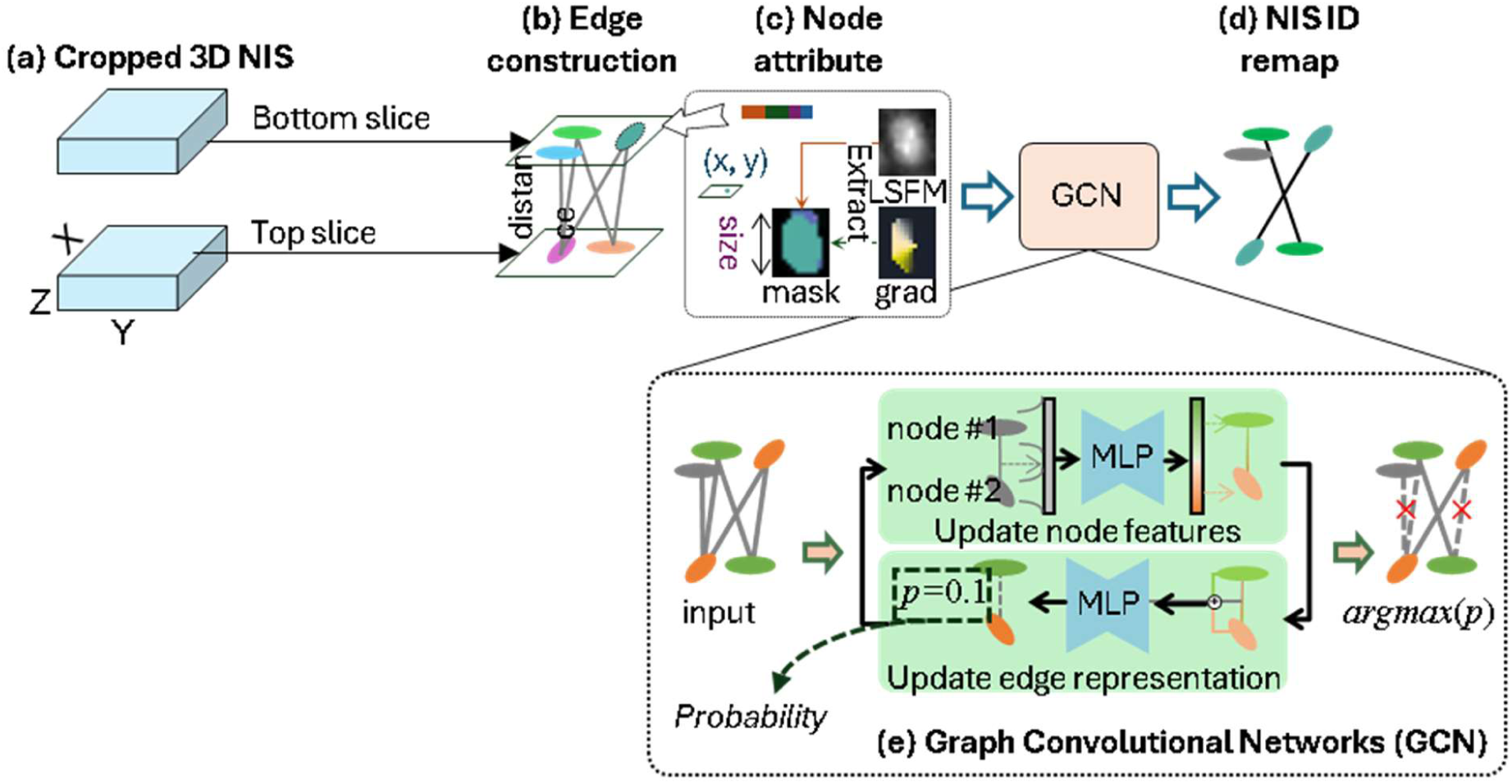
Learning-based stitching in the cropped area. a: Graph data construction for cropped segmentation masks in the third dimension (Z-dim). b: The edge is constructed by thresholding distance in the XY plane. c: The node attribute contains four elements: LSFM intensity, 3D gradient histogram denoted by ‘grad’, NIS size, and NIS location (x, y). d: The ID of matched NIS is remapped as the same according to the final graph. e: The architecture of graph convolutional networks (GCN), where ‘MLP’ denotes a multi-layer perceptron that consists of two layers of linear transformation using torch.nn.Linear with a ReLU activation in-between, and edges other than the max predicted *p* for each NIS mask in the top slice is removed.

**Extended Fig. 4:**
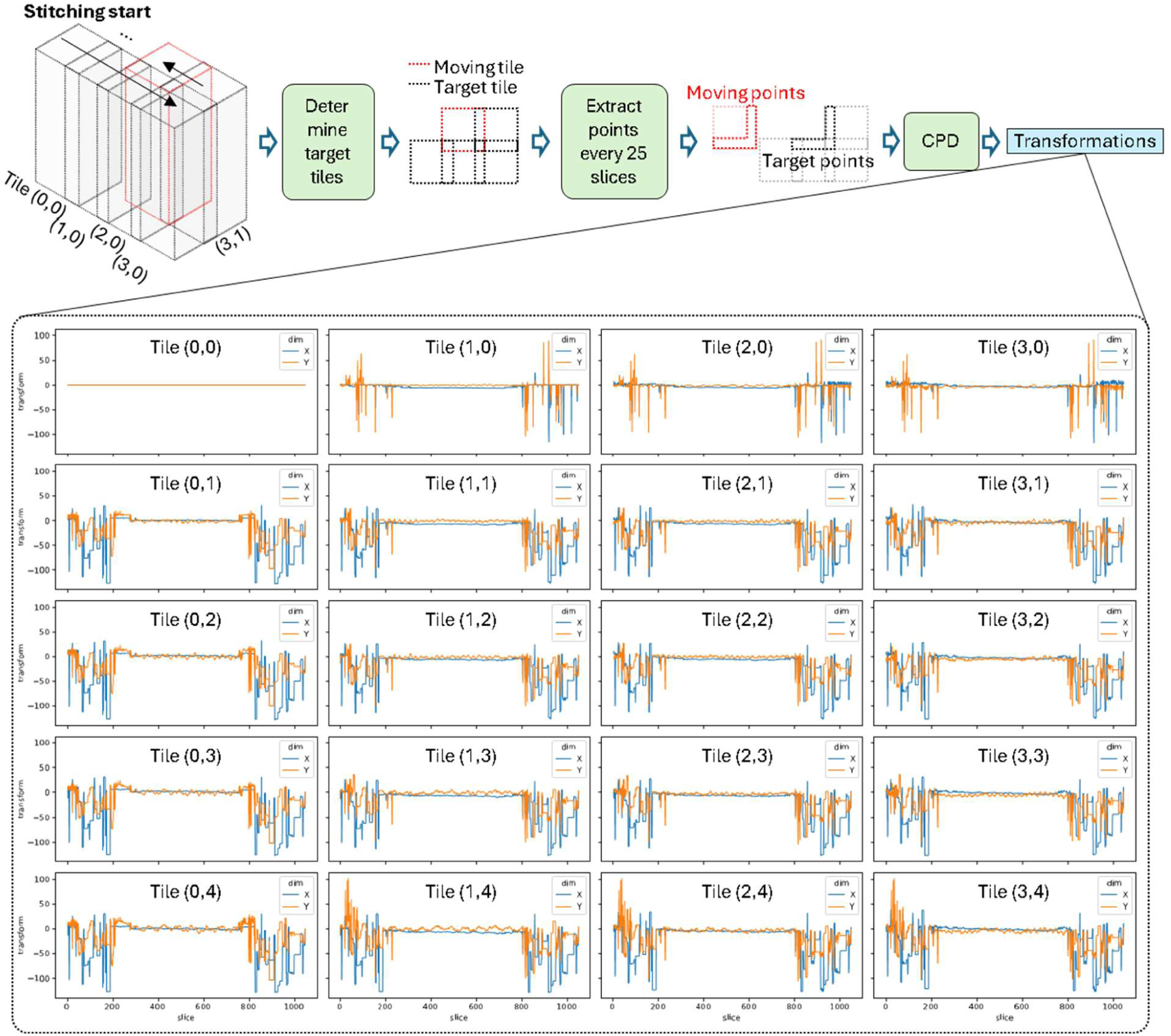
Automatic stitching for overlapping area of tiles in the XY plane. ‘CPD’ denotes the rigid registration version of Coherent Point Drift. We demonstrate the transformations of an example brain, where vibrations occurred during imaging for some top and bottom slices, and the vibration pattern is consistent within the same row.

**Extended Fig. 5:**
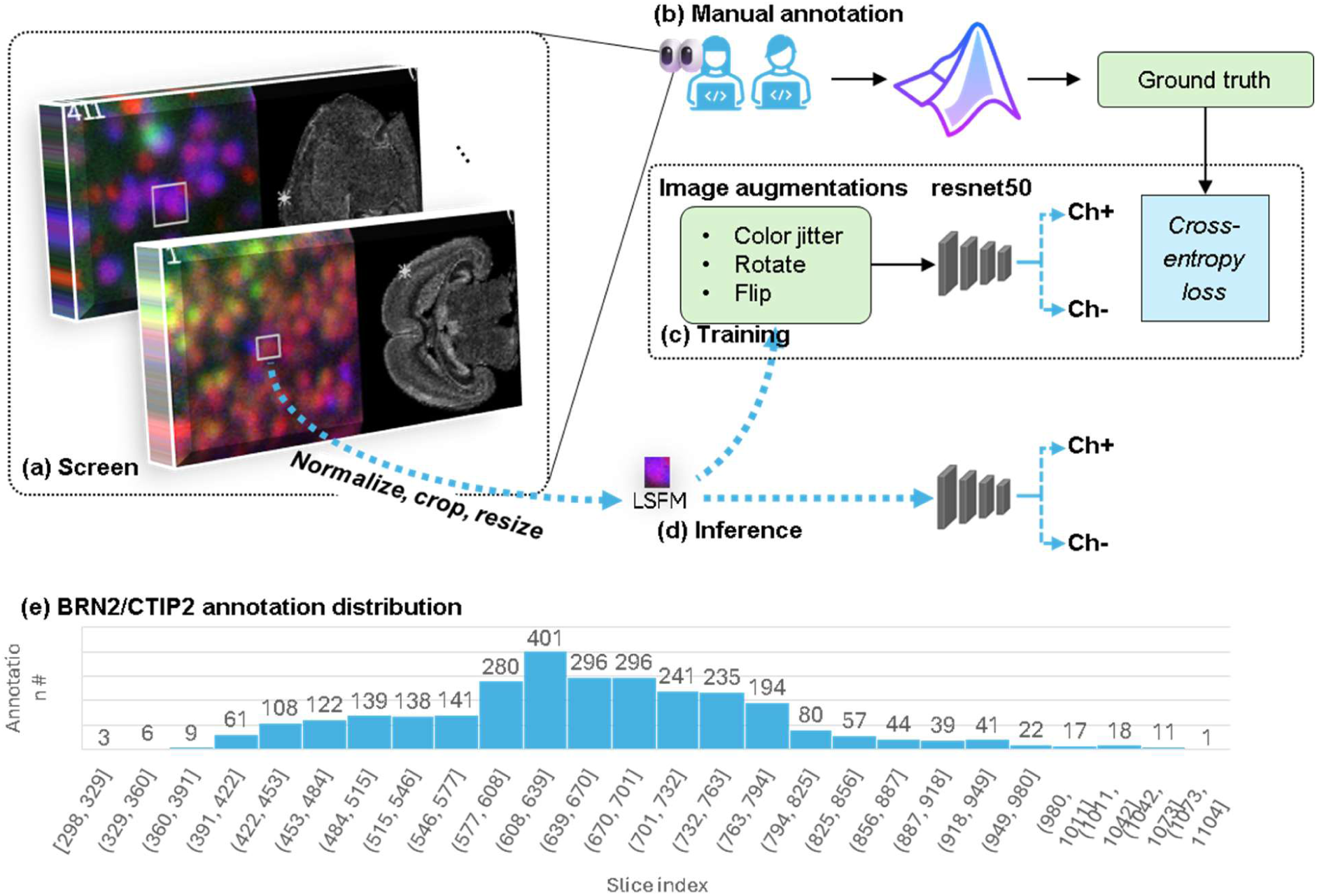
Learning-based multichannel co-localization method. a: Neuroscientists reviewing local patch images (100×100 pixels) with information of the global location marked by a white star over the whole-brain density map. The patches are examples of Brn2/Ctip2 and SOX9/NEUN, respectively. b: Neuroscientists use an interactive MATLAB tool to record if the detected cell is positive on a channel. c: The labeled multichannel patches are normalized, cropped, and resized based on the NIS bounding box for training a ResNet50 with 20% validating data. d: The best model weights are used for the inference of whole-brain co-localization.

**Extended Fig. 6:**
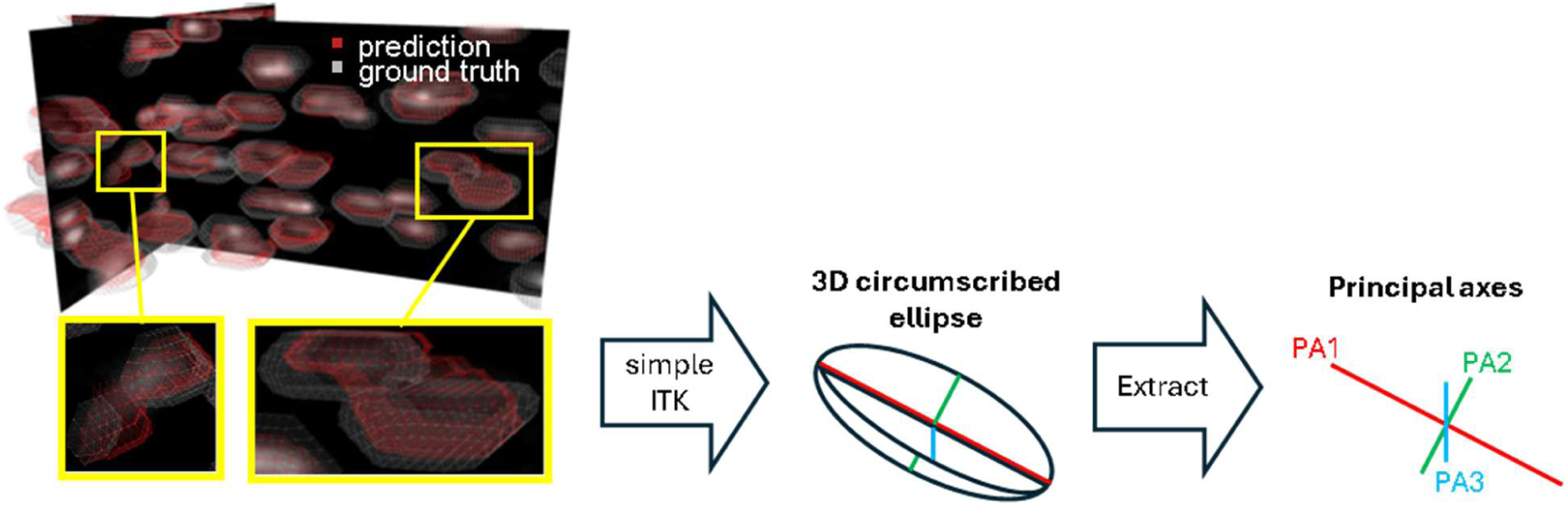
Cellular morphometry upon 3D NIS results of *CellPheno*. The simpleITK estimates three principal axes of each single instance segmentation mask to measure the ellipsoid, which is indicated by the length ratio between the second (PA2), the third (PA3), and the first principal axes (PA1).

**Extended Fig. 7:**
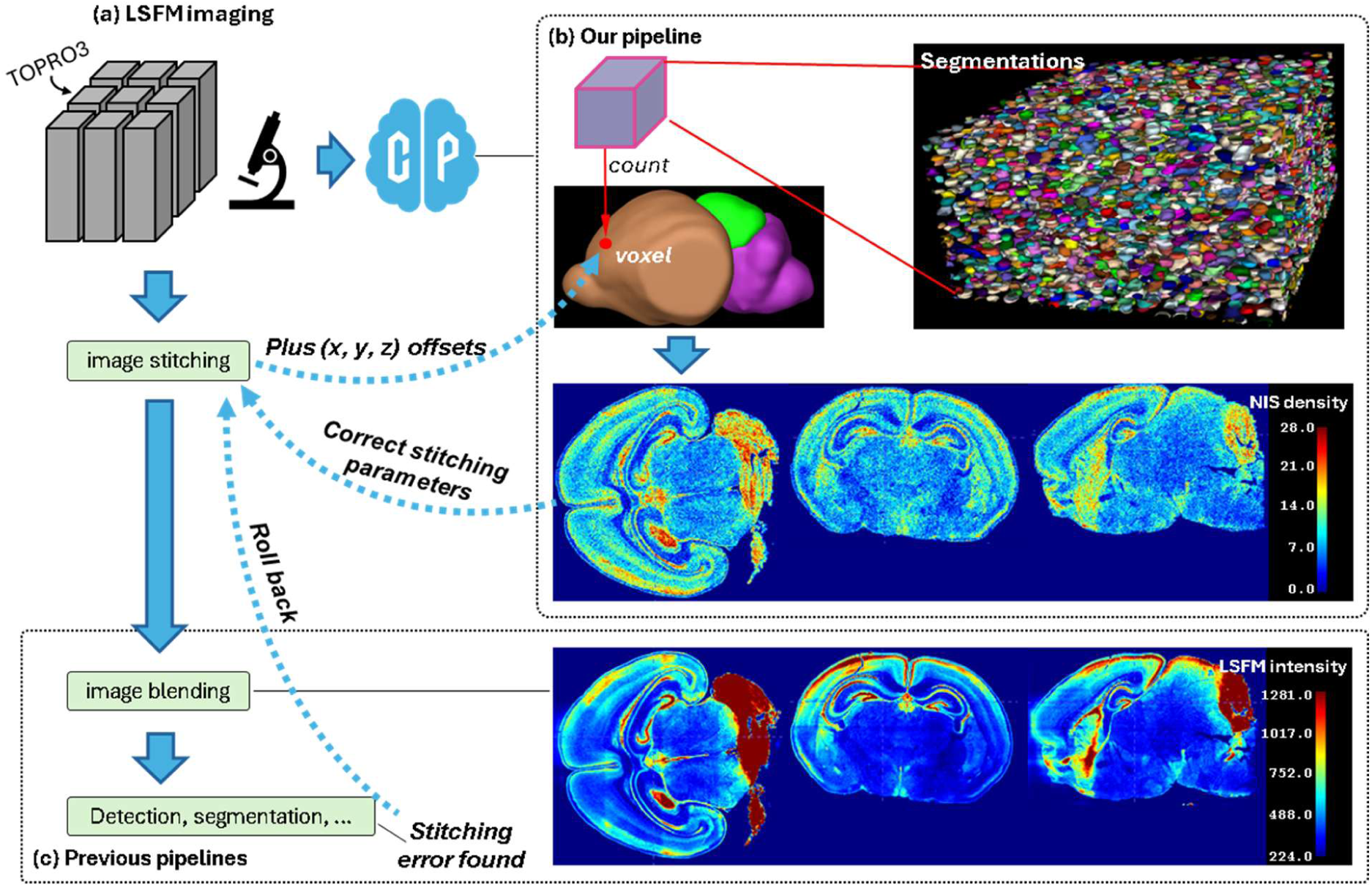
The *CellPheno* density map usage. a: Image stitching and blending are challenges since LSFM imaging is done tile-by-tile. b: Logic behind our pipeline is rooted in an independent whole-brain density map generation following those preprocessing steps. c: Since image stitching and blending in previous pipelines, taking NuMorph as an example, causes artifacts that are hard to find in downsampled LSFM images. Rolling back leads to a higher cost when a stitching error is found after image analysis.

**Extended Fig. 8:**
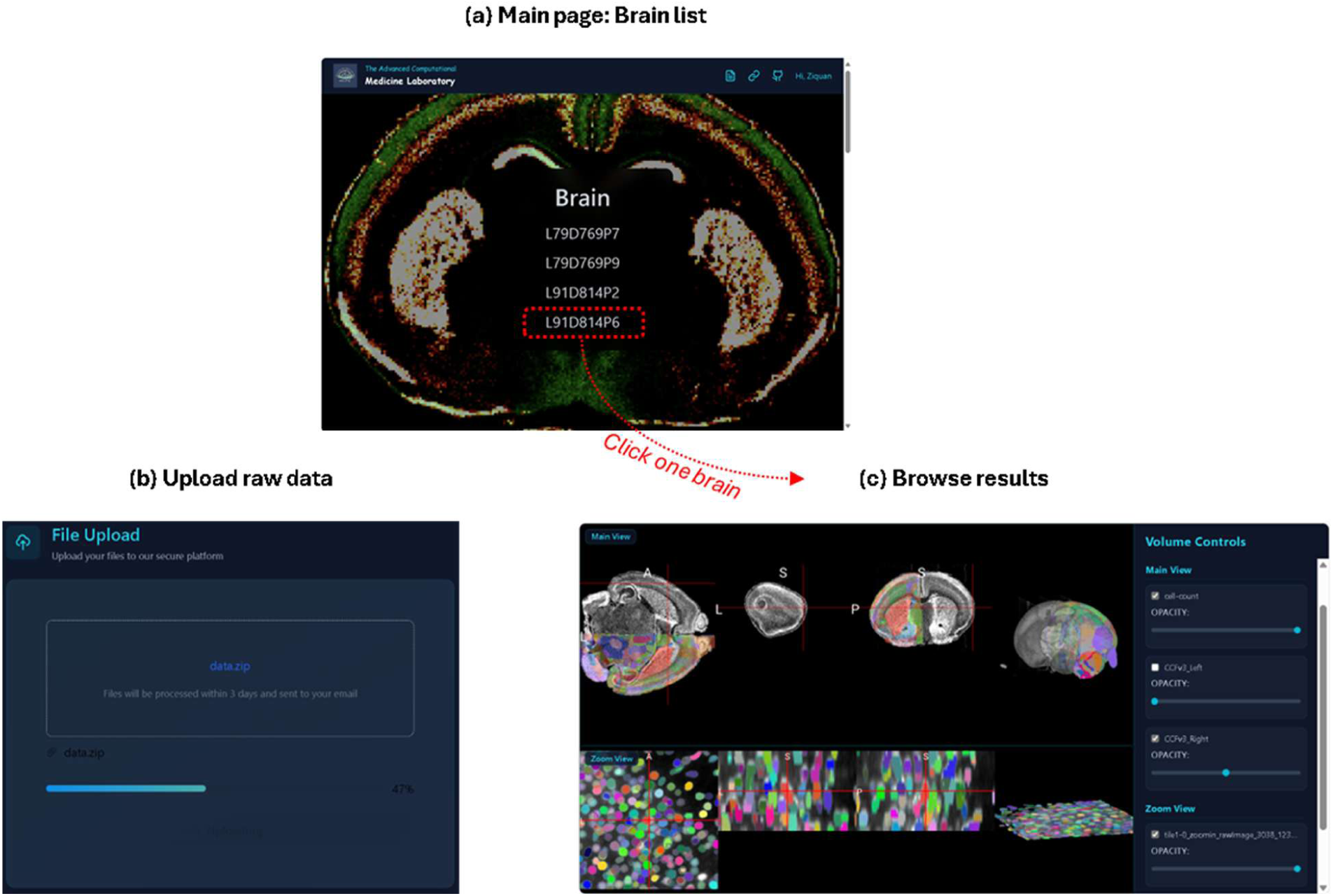
The website interface. a: The list of brains that are ready for browsing. b: Uploading raw data to the server. c: 3D NIS results browsing with whole-brain visualization (upper row) and the local zoom-in view (lower row).

