## Supplementary figures and images for "CellPheno: A High-throughput Computational Platform for Quantifying Cellular Resolution Whole Brain Microscopy Images"

### Extended Fig. 1

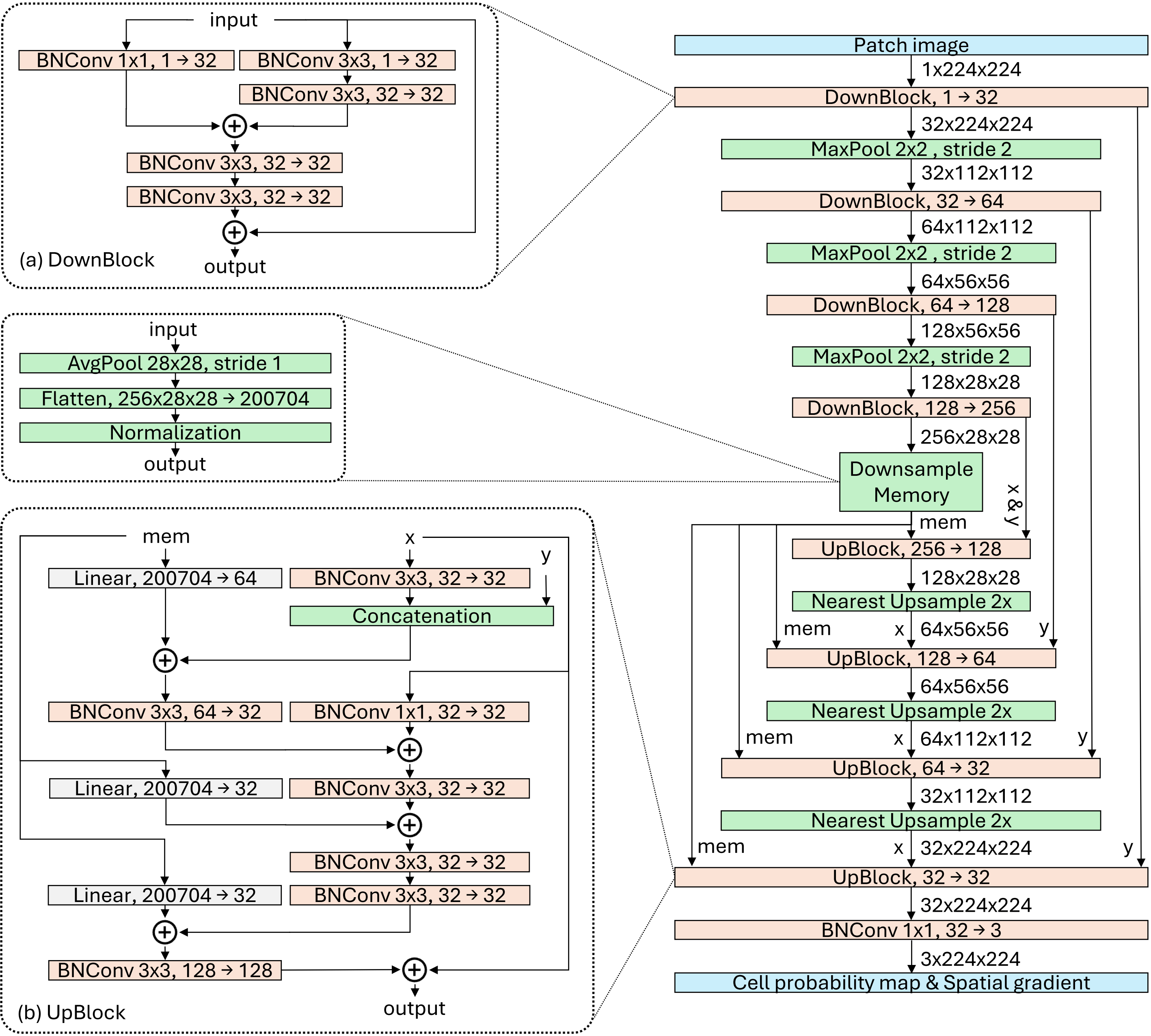

### Extended Fig. 2

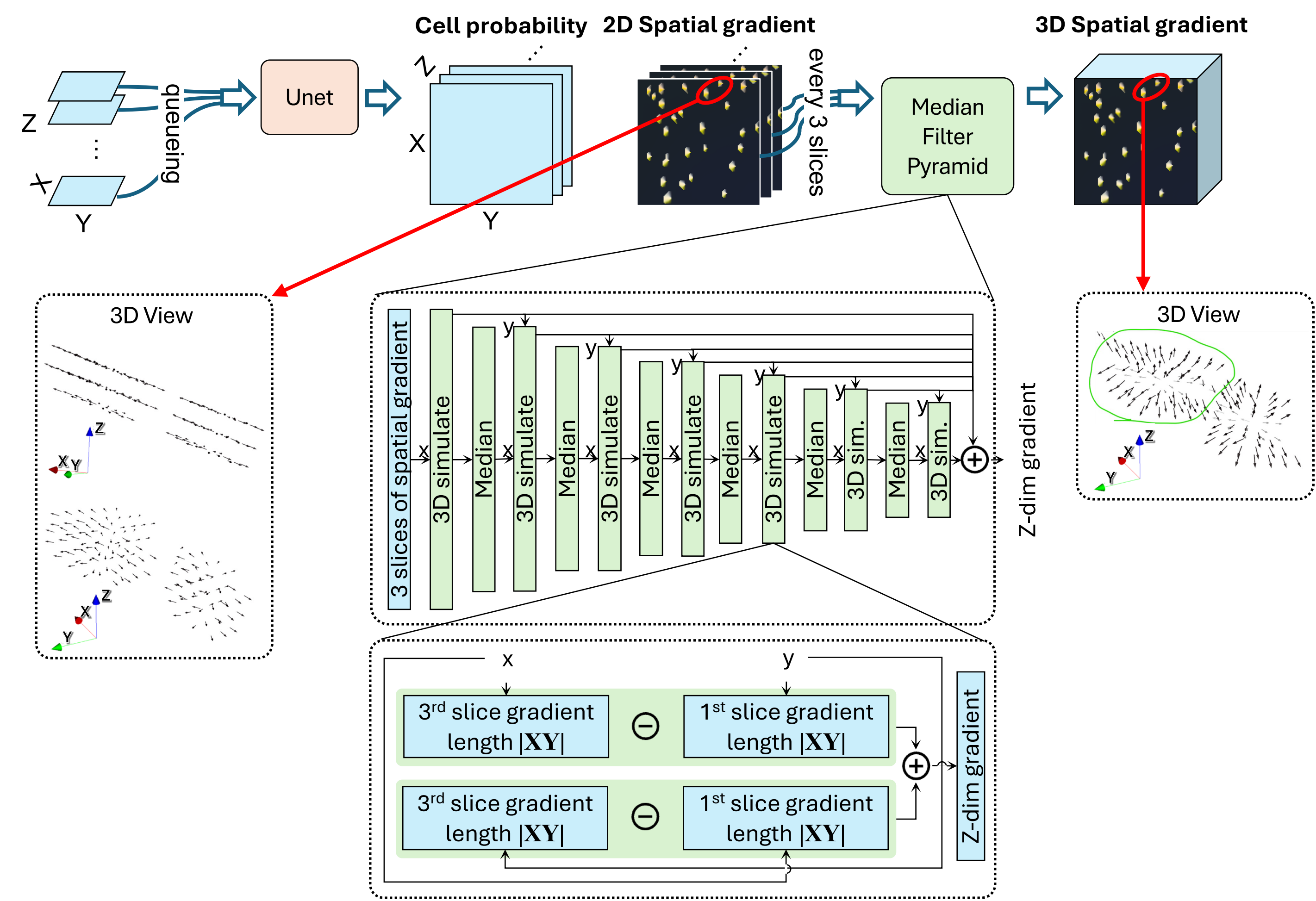

### Extended Fig. 3

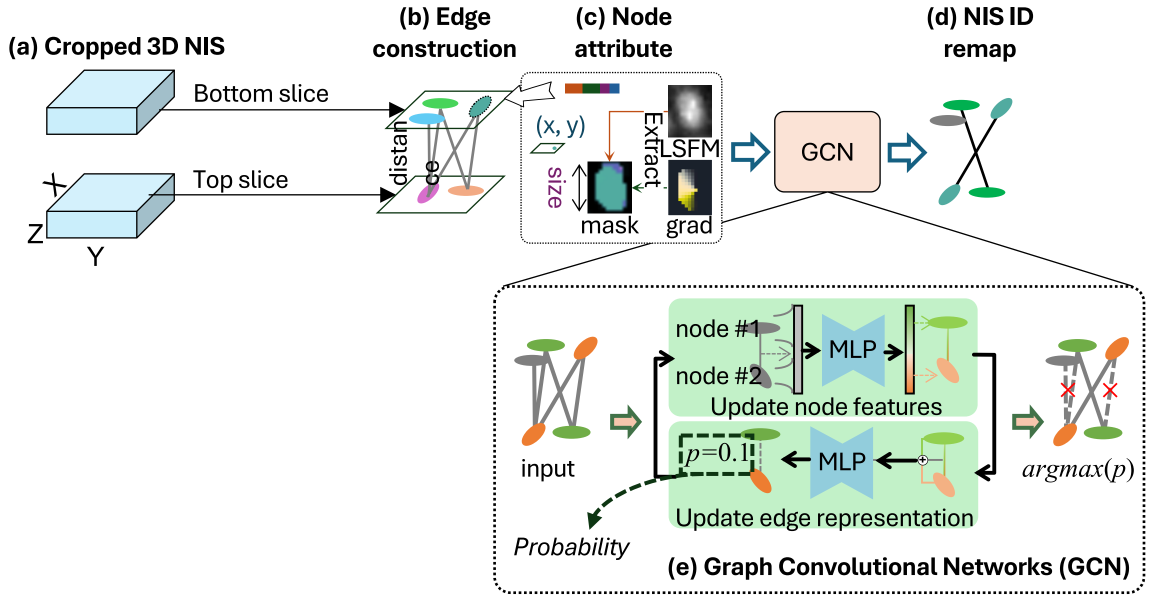

### Extended Fig. 4

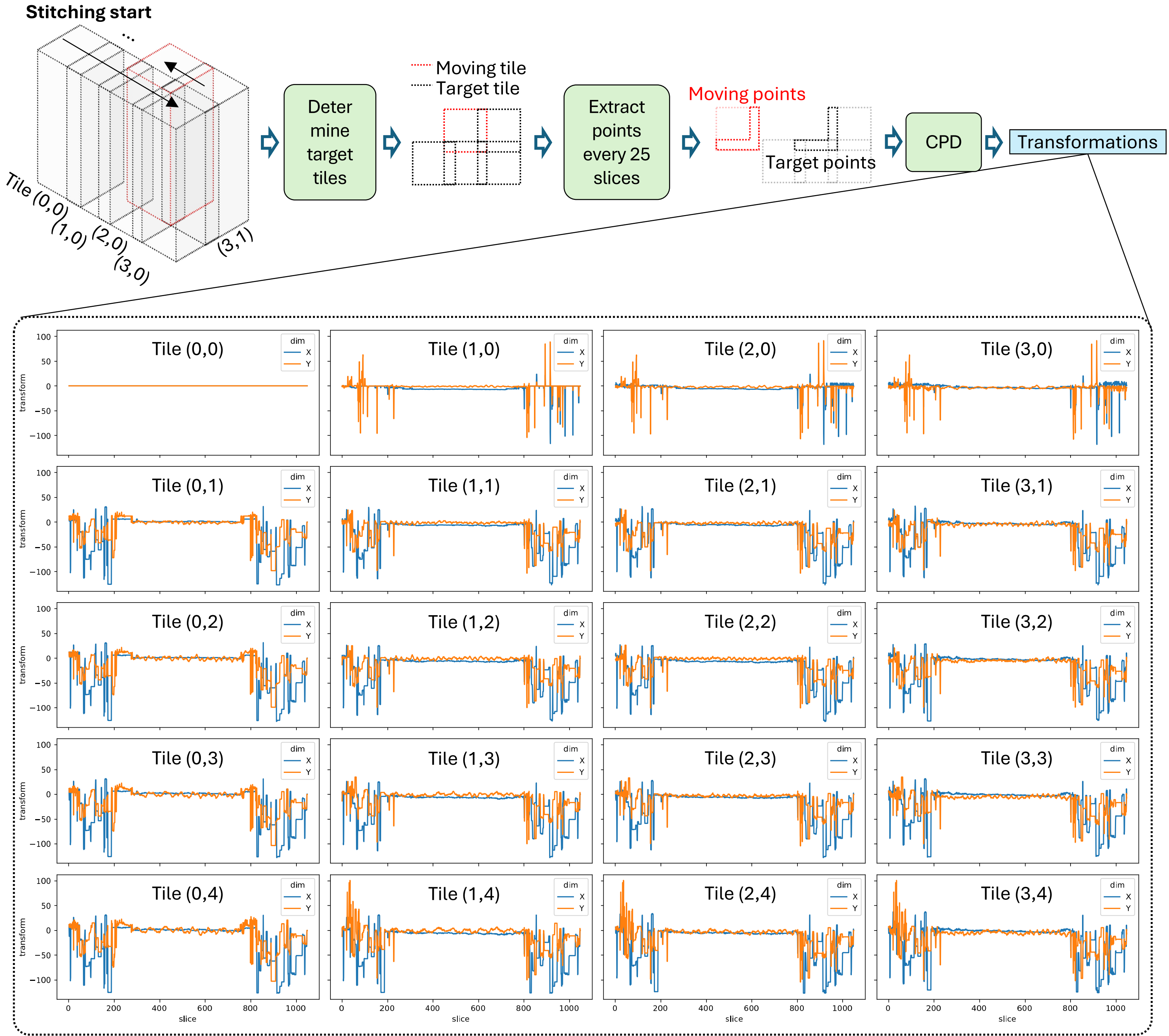

### Extended Fig. 5

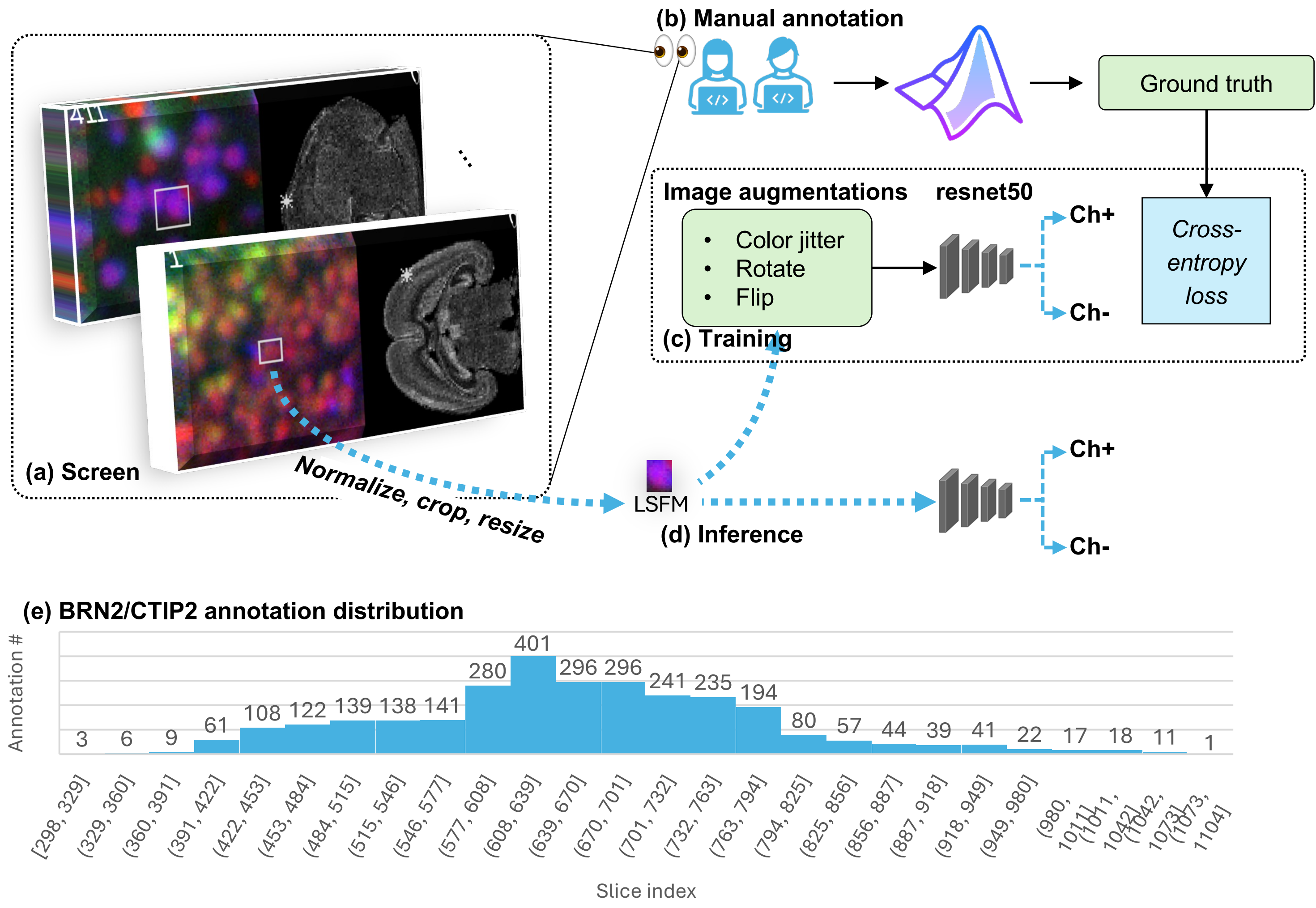

### Extended Fig. 6

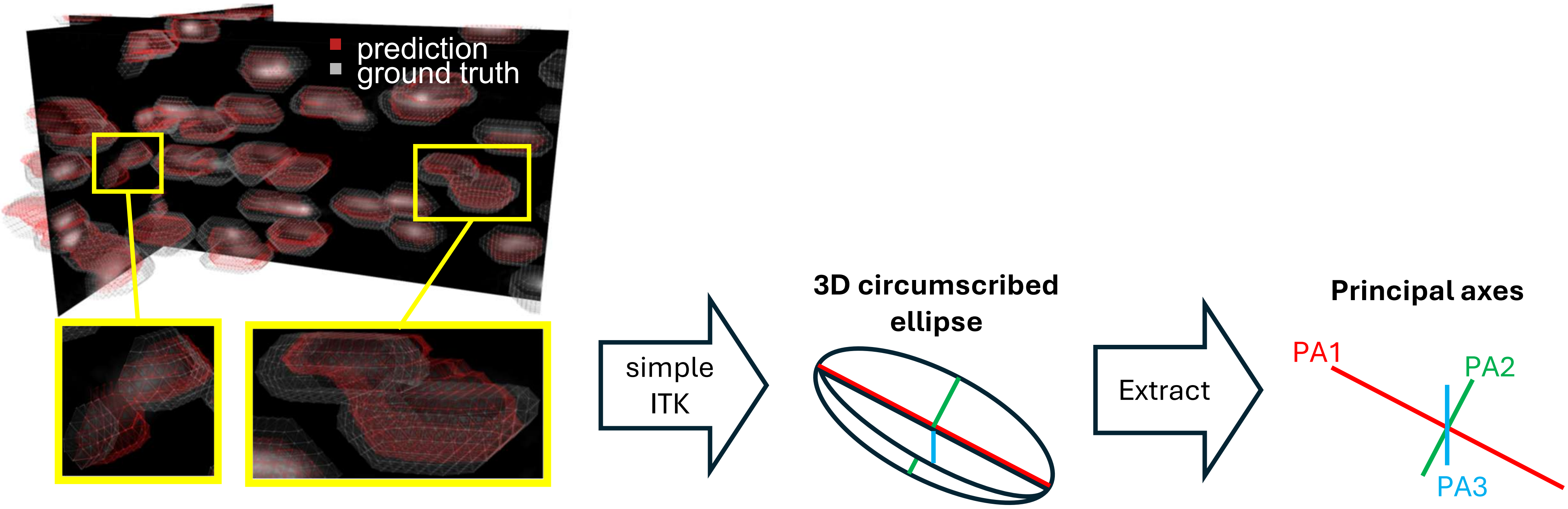

### Extended Fig. 7

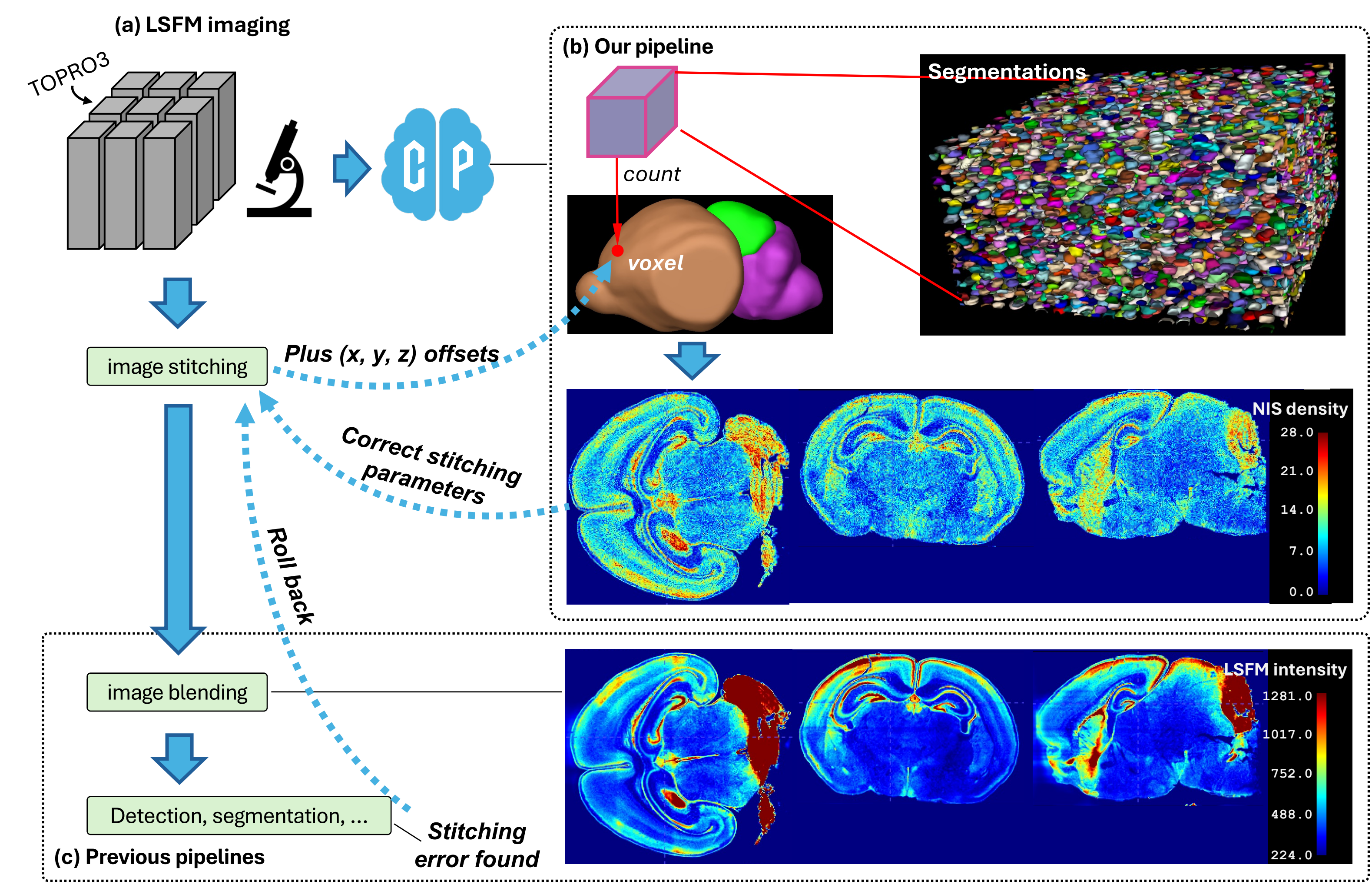

### Extended Fig. 8

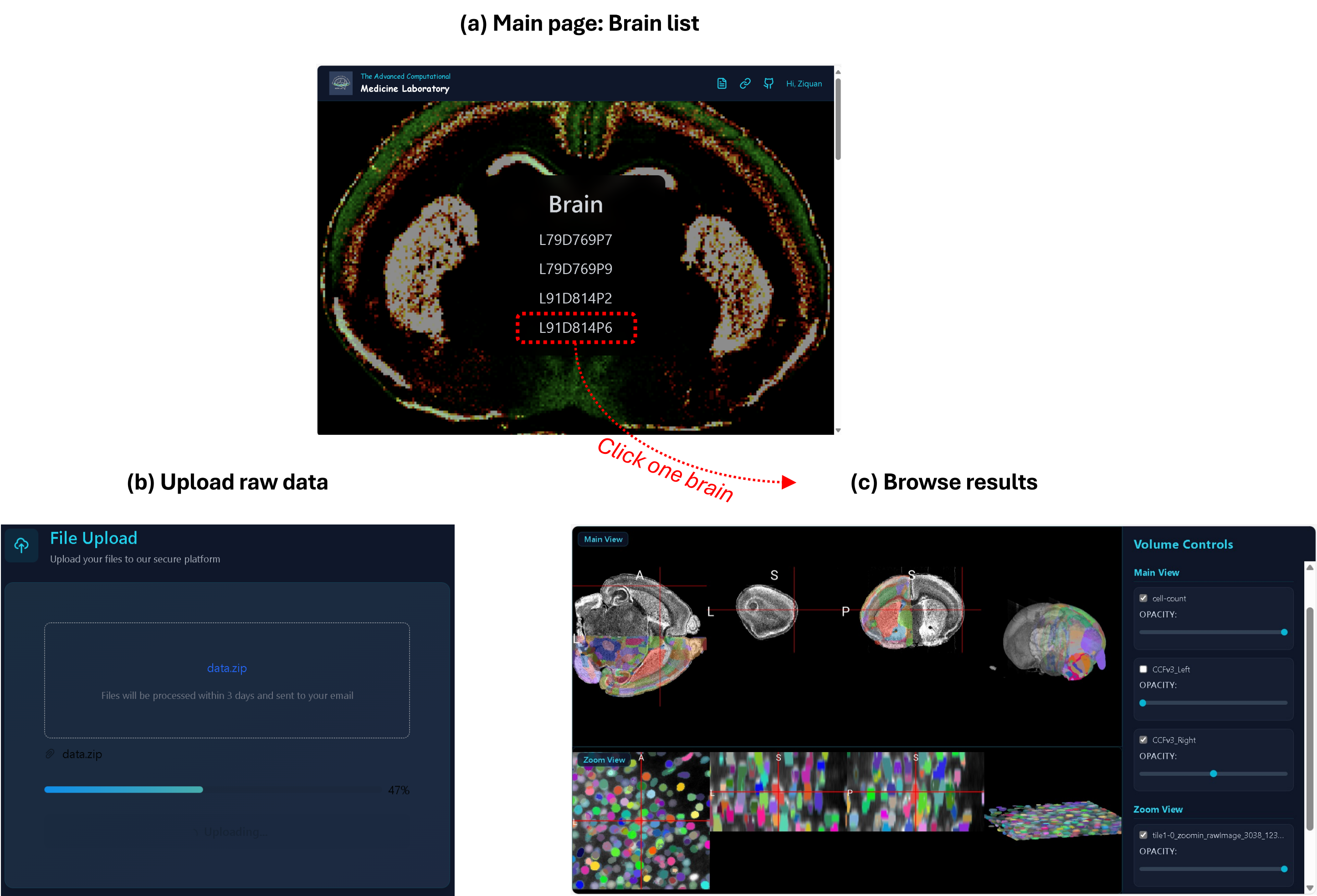
